# An inducible germ cell ablation chicken model for high-grade germline chimeras

**DOI:** 10.1101/2023.06.14.544953

**Authors:** Yi-Chen Chen, Daisuke Saito, Takayuki Suzuki, Tatsuya Takemoto

## Abstract

Chicken embryos are a powerful and widely used animal model in developmental biology studies. After the development of CRISPR technology, gene-edited chickens have been generated by transferring primordial germ cells (PGCs) after genetic modifications. However, the low inheritance caused by the competition between host germ cells and the transferred ones is the most common complication and largely reduces the production efficiency in this way. Here, we generated a gene-edited chicken, in which germ cells can be ablated in a drug-dependent manner, as recipients for gene-edited PGC transfer. We used the nitroreductase/metronidazole (NTR/Mtz) system for cell ablation, in which NTR produces cytotoxic alkylating agents from administered Mtz, causing cell apoptosis. The chicken Vasa homolog (CVH) gene locus is used to drive the expression of the NTR gene in a germ cell-specific manner. In addition, a fluorescent protein gene, mCherry, was also placed in the CVH locus to visualize the PGCs. We named this system **g**erm cell-**S**pecific **A**utono**M**o**U**s **R**emov**A**l **I**nduction (gSAMURAI). gSAMURAI chickens will be an ideal recipient to produce offspring derived from transplanted exogenous germ cells.

## Introduction

Chickens, as an oviparity organism, show an almost comprehensive embryonic development process *in ovo* and a comparatively accessible incubation condition compared to other animals, making them an ideal model for research in embryology since the times of ancient Greece (Zagris, 2022). For example, the chicken model allows accessible utility in embryo manipulation for xenogeneic tissue grafting to generate quail-chick chimeras, thus contributing to deciphering the ontogeny of the neural crest and of the immune system (Le Douarin, 1988). In addition, with an ex ovo embryo culture system, namely, “New’s culture”, and embryo electroporation, chickens have become non-mammalian model animals due to advances in research techniques for studying gene regulation and function in development (New, 1955; Uchikawa et al., 2004). Although chicken embryos provide convenience for developmental biologists, the difficulty of transgenic chicken production has limited the merits of chicken utilization in developmental biology studies. Therefore, the availability of a simple method for the generation of transgenic chickens would expand the usefulness of chickens and lead to revolutions in understanding developmental regulation.

The production of transgenic chickens has been difficult and presents an inconvenience for the usage of research due to their unique reproductive system. In mammalian models, a 1-cell stage zygote is accessible and utilized to generate transgenic animals. However, eggs shed from the chicken oviduct have already developed to a high cell number, and a 1-cell stage zygote of the chicken emerges in the infundibulum of the reproductive tract and fertilization is polyspermy, indicating an inconvenience to produce transgenic chicken. Therefore, transgenesis in chickens has mainly depended on the viral transfection system (Sang, 1994). Another strategy to generate transgenic animals is to utilize embryonic stem cells. Although chicken embryonic stem cells (cESCs) have been established, none of the contributions to germline lineage and offspring production from cESCs have been reported (Lavial and Pain, 2010; Pain et al., 1996; van de Lavoir et al., 2006b). Therefore, the accessibility of gene-edited organisms between chickens and others presents a huge difference, particularly after the development of CRISPR technology.

Nevertheless, by using genetic modification in the germline, genetically modified chicken offspring are able to be obtained (Salter et al., 1987). For better efficiency of germline transmission, scientists have focused on the improvement of in vitro culture methods and genetic manipulations of chicken primordial germ cells (PGCs) (van de Lavoir et al., 2006a). With many of their contributions, this PGC-mediated gene-editing technique has become widely used not only in basic scientific research to identify the mechanism of germ cell development or sexual determination but also in industrial purposes, such as the generation of transgenic chickens for recombinant protein production and breeding for low-allergen chickens (Ching et al., 2018; Choi et al., 2022; Ioannidis et al., 2021; Oishi et al., 2016).

However, the major disadvantage of this method is the low germline transmission efficiency caused by the competition between recipient host germ cells and donor cells. In addition, it is known that the efficiency becomes lower when chicken PGCs undergo long-term in vitro culture or manipulation (Collarini et al., 2015; Woodcock et al., 2019). To solve this problem, the chemical or irradiation method was adopted to remove the endogenous germ cells of the recipient embryo before PGC transfer, while higher mortality in the treated embryo was found as the major side effect (Nakamura et al., 2012; Nakamura et al., 2010).

In this study, we established a new chicken model to be used as a recipient chicken for PGC transplantation, in which host germ cells can be removed in a drug-dependent manner. We utilized the nitroreductase/metronidazole (NTR/Mtz) system for cell ablation, in which NTR produces cytotoxic alkylating agents from administered Mtz, causing cell apoptosis (Chen et al., 2011; Curado et al., 2008).

## Materials and methods

### Chicken and chicken embryos

For PGC donor embryos, Barred Plymouth Rock (BPR) strain chicken (*Gallus gallus*) fertile eggs were purchased from Okazaki station, National Livestock Breeding Center, Japan. For recipient embryos of gSAMURAI PGC transfer, fertile eggs of JuliaLite strain chickens (*Gallus gallus*) were purchased from Japan Layer K.K., Japan. gSAMURAI fertile chicken eggs were obtained by artificial insemination at Kyodoken Institute, Japan, or by purchasing from Okazaki station, National Livestock Breeding Center, Japan. All animal procedures were approved by the Institutional Animal Care and Use Committee of Tokushima University (T2021-53) and Kyodoken Institute (A2021-001-3).

### Plasmid construction

The ST7 expression plasmid was synthesized by using the public sequence of the pX330-U6-Chimeric_BB-CBh-hSpCas9 plasmid (Addgene #42230)(Cong et al., 2013) as a backbone and replacing the regions for a Cas9 CRISPR RNA (crRNA) and a fragment from 3xFLAG to nucleoplasm NLS with a MAD7 crRNA and a cassette containing a C-terminal NLS-conjugated MAD7 nuclease (nMAD7), respectively. The P2A-mScarlet fragment was obtained by PCR cloning from pEB2-mScarlet (a kind gift from Dr. Philippe Cluzel; Addgene #104006)(Balleza et al., 2018) and inserted following the 3xHA by assembling with NEBuilder HiFi DNA Assembly (New England Biolabs, USA). The donor plasmid for homologous recombination was generated by inserting a synthetic cassette containing T2A-NTR-T2A-Puromycin N-acetyltransferase-T2A-mCherry at the location before the stop codon of the CVH coding DNA sequence (CDS) with NEBuilder HiFi DNA Assembly. The homology-arm sequence was cloned in a genomic DNA (gDNA) template (chromosome Z: 17501775-17503792, bGalGal1.mat.broiler. GRCg7b) and inserted into the pBluescript II SK(+) plasmid by TA cloning. All synthetic DNA fragments were purchased from Integrated DNA Technologies Inc. (USA).

For crRNA cloning, a pair of BbsI enzymatic digestion sites were designed to set just after the MAD7 crRNA in the ST7 expression plasmid for the cloning of annealed crRNA oligos by restriction cloning with BbsI and T4 ligase (New England Biolabs, USA). crRNA candidates were designed with CHOPCHOP (Labun et al., 2019), and oligos were synthesized and purchased from Eurofins Japan (Japan).

### Cell culture and manipulation

Donor PGC clones were derived from E3 BPR strain chick embryos and maintained with the method described by Chen et al. (2019). The derived PGC clones were stored at -80 °C within Bambanker freezing medium (NIPPON Genetics, Japan) for further usage. The culture medium for PGC was composed of a basic medium and supplements (Table S1). For genetic modification in PGCs, one microgram of plasmid DNA was used for 5×10^4^ cells in a 10 μL electroporation reaction. For KI purposes, the ratio of ST7 expression plasmid and donor plasmid was 1:2 (w/w). Electroporation was conducted with an optimized parameter of a 1150 V/30 ms/1 pulse with buffer R in a Neon Transfection System (Thermo Fisher Scientific, USA). The PGC culture medium in the presence of puromycin (0.1 μg/mL; Sigma-Aldrich, USA) but no other antibiotics was used for puromycin selection (Chen et al., 2018).

### FACS sorting and analysis

For fluorescent cell enrichment sorting and analysis, experiments were conducted by a BD FACSMelody cell sorter (BD Biosciences, USA) with the default configuration and equipped with blue (488 nm), red (640 nm), yellow green (561 nm) lasers and BD Chorus software (BD Biosciences, USA). The instrument operation was performed according to the instructions in the user’s guide.

### DNA extraction and amplicon sequence analysis

gDNA samples from cells were prepared with a DNeasy Blood & Tissue Kit (Qiagen, USA). The DNA fragments around the crRNA target site were amplified by a two-step PCR with specific primer sets (Table S2) and Index PCR Primers mentioned in the manufacturer’s instructions (Illumina, USA). After gel purification, the amplicons were subjected to MiSeq for amplicon sequence analysis by using the MiSeq Reagent Kit v2 (Illumina, USA).

### Immunofluorescence staining

Cells were attached to a hydrophilic micro slide glass (Matsunami Glass, Japan) and adapted to immunofluorescence staining with the method described previously (Chen *et al*., 2019). Cells were labeled with anti-DAZL (200×; Abcam, UK), anti-DDX4 (200×; Bioss, USA), anti-NANOG (1000×; a kind gift from Dr. Kiyokazu Agata)(Nakanoh et al., 2015), and anti-mCherry (500×; Novus Biologicals, USA) antibodies and then subsequently stained with Alexa Fluor™ 488-conjugated donkey anti-rabbit IgG (2000×; Invitrogen, USA) and Alexa Fluor™ 555-conjugated donkey anti-mouse IgG (2000×; Invitrogen, USA) for visualization. Hoechst 33342 (2.5 μg/mL; Molecular Probes, USA) was used to stain the nucleus. Images were acquired using a Leica DMI4000 B microscope (Leica Microsystems, Germany) equipped with a digital CMOS camera (Hamamatsu Corporation, Japan).

### Chicken embryo culture and dissection

In ovo chicken embryos were incubated with a humidified egg incubator (Showafuranki, Japan) at 38 °C, and automatic egg turning was performed. For ex ovo embryo culture, E2.5 embryos were excised from the yolk and cultured on a petri dish with a method described by Uchikawa *et al*. (2004) for observation. For embryo dissection, the embryo was isolated at E10, E12, or E16. Gonads were derived by gently picking them up with tweezers and then washing once in cold PBS without Ca2+/Mg2+ (Takara, Japan). Image and movie acquisition was conducted by a Leica M205 FA fluorescence histological microscope (Leica Microsystems, Germany) equipped with cellSens Standard (Olympus, Japan).

To prepare the sample for FACS analysis, gonad tissue was dispersed in TrypLE™ Express Enzyme (Gibco, USA) with gentle shaking for 10 min. After centrifugation (600 g, 5 min), dispersed tissue was resuspended in PBS without Ca2+/Mg2+ and passed through a 100 μm strainer (Falcon, USA) for further analysis.

### PGC transplantation and prodrug supplementation

A total of 5×10^4^ PGCs were transplanted into the circulation of E3 recipient in 5 μL of a basic medium or prodrug-containing medium with the method mentioned by Chen *et al*. (2019) for gonad migration assay. Prodrug-containing groups were prepared just before the transplantation by diluting some volume of cell suspension and the basic medium of Mtz presence in 2 mM or 10 mM to meet the final concentration in 1 mM or 5 mM, respectively. Mtz was purchased from Fujifilm Wako Pure Chemical Corporation (Japan). Data were analyzed with GraphPad Prism9 (GraphPad Software, Inc., USA) using one-way analysis of variance (ANOVA) and Fisher’s least significant difference (LSD) test. Differences between groups were considered to be significant at a *P* value of <0.05 or 0.01.

## Results

### Design of a **g**erm cell-**S**pecific **A**utonoMoU **R**emov**A**l **I**nduction (gSAMURAI) chicken

To remove endogenous germ cells in a drug-dependent manner, we applied the nitroreductase/metronidazole (NTR/Mtz) system (Curado *et al*., 2008). We inserted a bacterial nitroreductase (NTR) gene into the 3’ end of the chicken Vasa homolog (CVH) coding sequence via the 2A peptide sequence using gene editing-mediated homologous recombination. The puromycin resistant (puro) gene for the selection of recombinants in PGCs and the mCherry gene for the visualization of PGCs were also inserted into the same locus (Fig. 1).

**Fig. 1.**
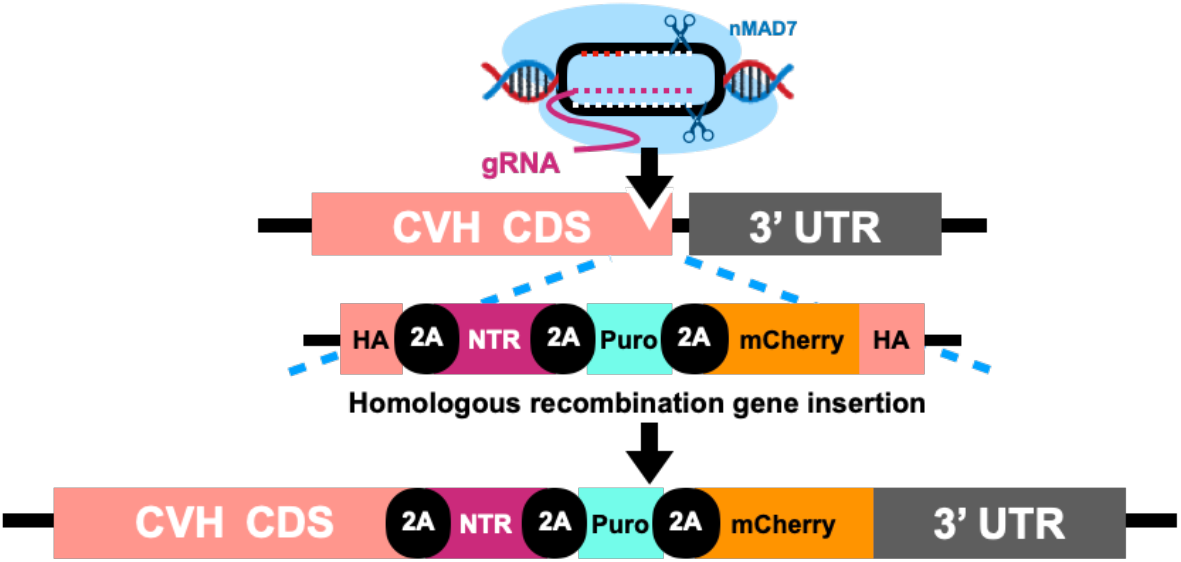
Schematic of gSAMURAI chicken

### Selection of crRNA of MAD7 nuclease for optimized gene editing efficiency

The higher the double-stranded break (DSB) efficiency, the higher the knock-in (KI) efficiency. To this end, we first identified the optimized crRNA to introduce mutations into the CVH gene using chicken PGCs. Three crRNAs targeting the 3’ end of the CVH gene were designed, and their insertion–deletion mutation (Indel) rates, which reflect DSB efficiency, were examined (Fig. 2A). We constructed a plasmid expressing each crRNA, MAD7 nuclease plus a C-terminal NLS (nMAD7), and mScarlet ubiquitously using U6 or CBh promoters. Each plasmid was introduced into chicken PGCs by electroporation. PGCs were derived from blood vessels of embryonic Day 3 (E3) Barred Plymouth Rock (BPR) strain embryos, undergoing approximately 3 weeks of in vitro propagation until the cell number was sufficient for further studies. At Day 2 posttransfection, successfully transfected cells were collected through FACS sorting by positive expression of mScarlet (Fig. 2B and 2C), and genomic DNA was extracted for indel analysis at each target site. Sequence analysis of genomic PCR amplicons revealed that among these three crRNAs, No. 3 had the highest indel frequency at 9.59% (Indel read/Total read: 105/1059), compared to No. 1 and No. 2 crRNAs, which presented 1.99% (Indel read/Total read: 20/1003) and 1.32% (Indel read/Total read: 14/1064), respectively. We used crRNA NO.3 (Fig. 2D) for further experiments.

**Fig. 2.**
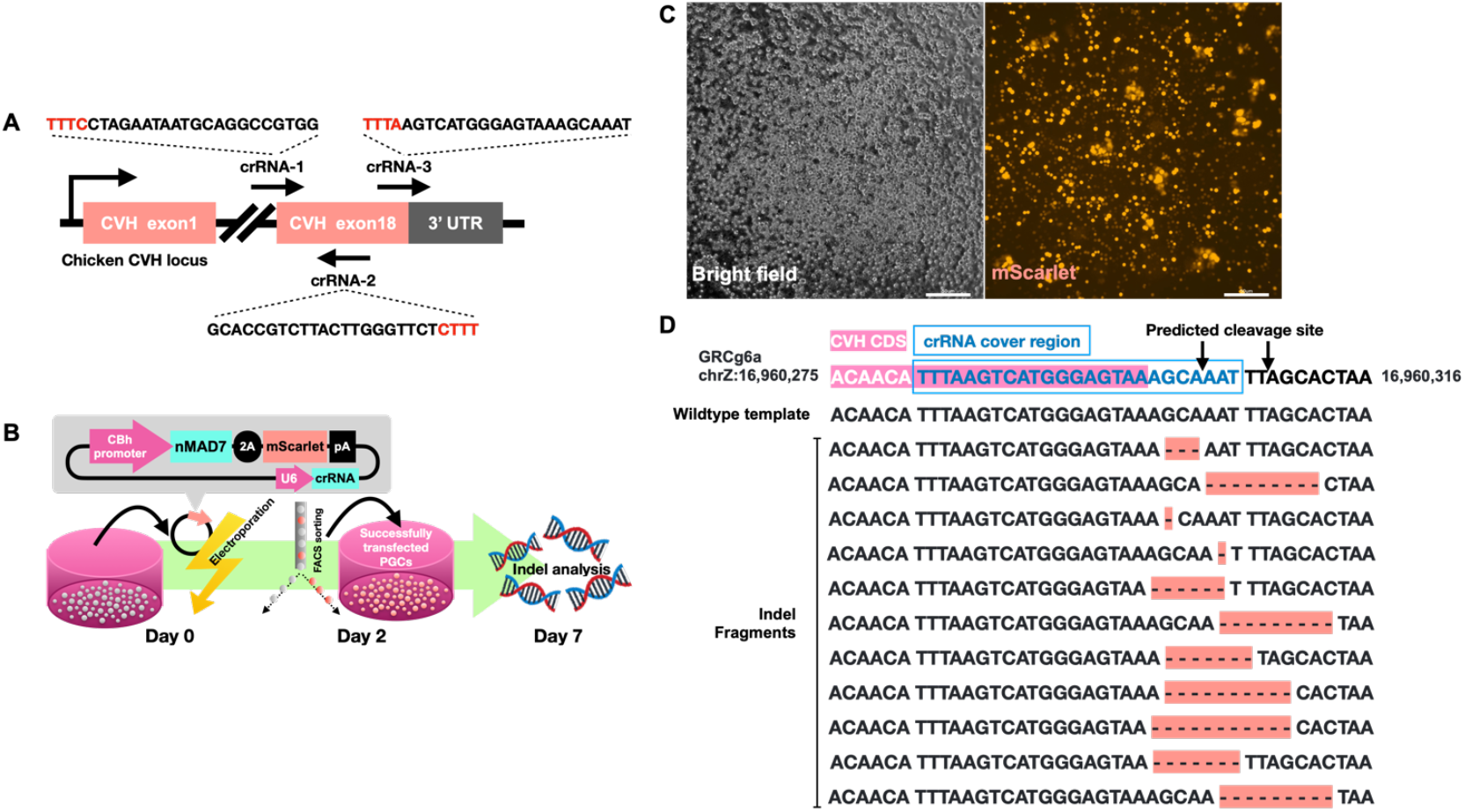
Evaluation of nMAD7 crRNAs showing high indel formation in chicken PGCs. (A) Three ST7 crRNAs targeting the C-terminus of the CVH coding region were designed. The recognition sequence of each ST7 crRNA is shown, and each PAM site is labeled in red. (B) Experimental procedure of plasmid transfection and the analysis of indel formation. First, the plasmid expressing ST7, crRNA and mScarlet was electroporated into PGCs. Second, successfully transfected cells were harvested by FACS sorting with positive mScarlet expression at Day 2 posttransfection. Finally, all sorted cells at Day 7 posttransfection were adapted to DNA extraction for further indel analysis by NGS. (C) mScarlet could be detected in PGCs 48 hours after electroporation (Scale bar: 50 μm). (D) NGS analysis data of PGCs electroporated with crRNA-3. Multinucleotide deletions around the predicted cleavage site (arrow) were found.

**Table 1.**
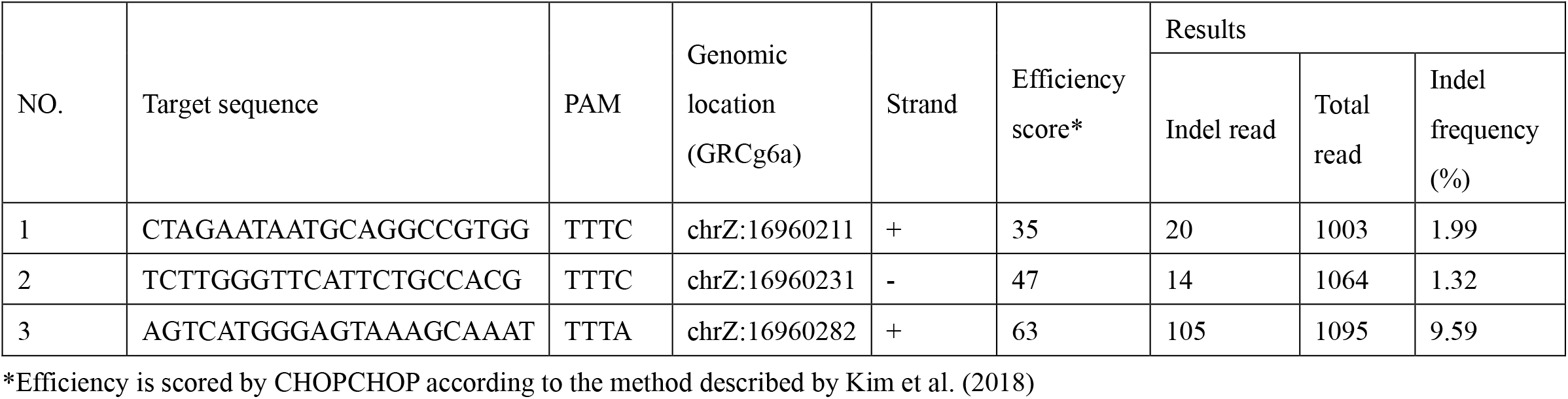
Designed crRNAs for CVH and their results of indel formation

### Establishment of PGCs carrying the gSAMURAI system by DNA cleavage-mediated homologous recombination

Using the nMAD7 expression plasmid and the donor plasmid, homologous recombinant PGCs were selected by mCherry fluorescence and puromycin resistance (Fig. 3A). As illustrated in Fig. 3B, mCherry-expressing PGCs were found in cell populations of both sexes in the second week after transfection at 0.53% (female) and 0.34% (male). After puromycin exposure for one week, the mCherry-expressing PGC rate was obviously enriched. By using a FACS sorter for quantification, 98.66% (female) and 99.39% (male) of mCherry-positive PGCs were detected after 2 weeks of antibiotic selection (Fig. 3B and 3C), indicating that the puromycin resistance gene supported the cell enrichment of KI PGCs. Moreover, two populations with different mCherry fluorescent intensities were found in the male PGCs, suggesting heterozygous and homozygous transgene insertion in CVH (Fig. 3C). To confirm successful knock-in, we performed genomic PCR analysis, followed by sequence analysis, using genomic DNA samples extracted from WT PGCs and PGCs with high mCherry intensity (Fig. 3D). The size and DNA sequences of each amplicon demonstrated that the transgene had been inserted into the targeted loci as in our design, gSAMURAI.

**Fig. 3.**
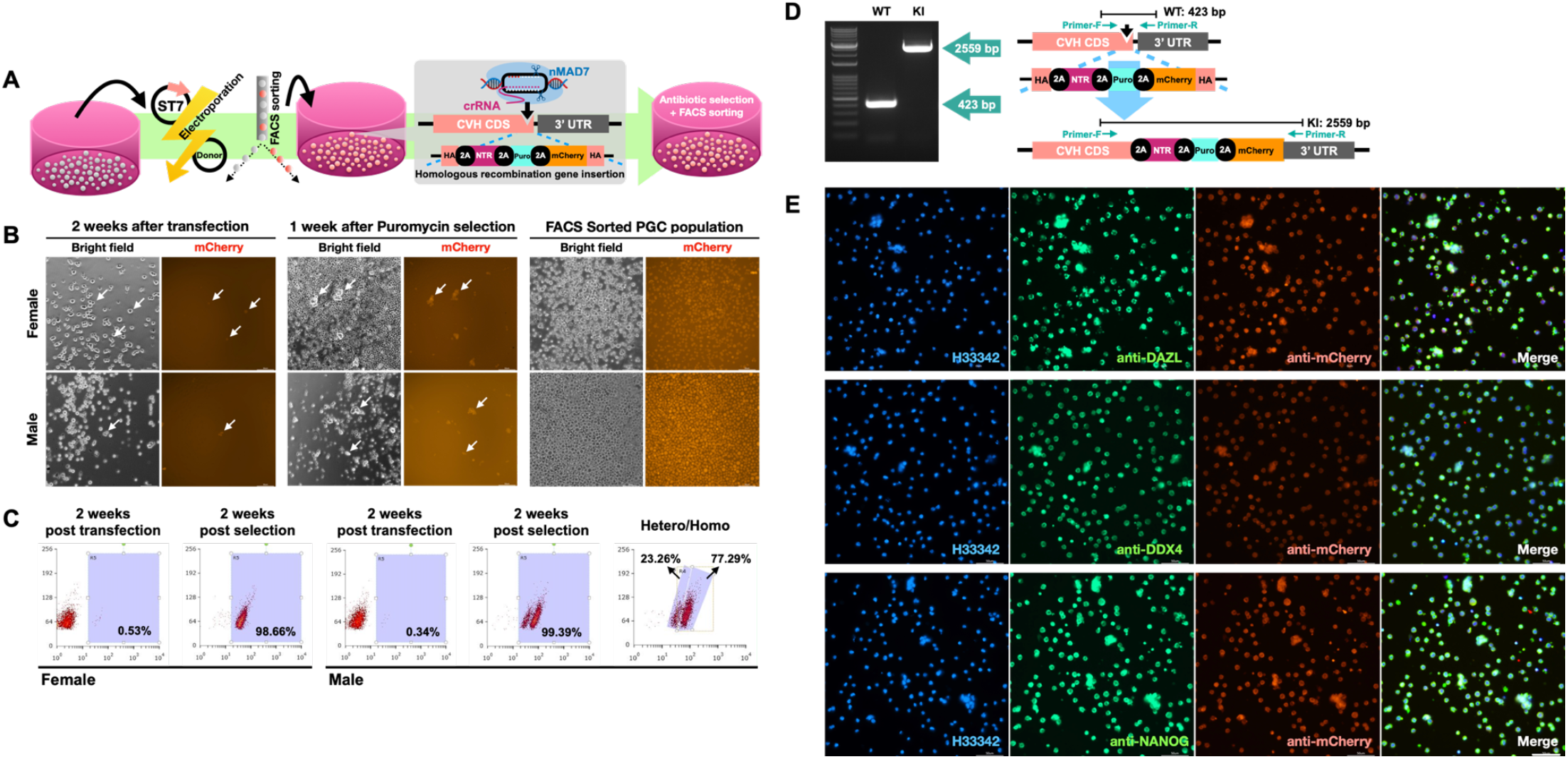
Generation of gSAMURAI PGCs utilizing DNA cleavage-mediated homologous recombination. (A) Illustration of the flowchart for the establishment of genetically modified PGCs through DNA cleavage-mediated homologous recombination. (B) The derivation and enrichment of genetically modified PGCs after transfection, antibiotic selection, and FACS sorting. (C) Cell population gating by the mCherry fluorescence intensity in PGCs after transfection and antibiotic selection. (D) PCR amplification using Primer-F and Primer-R for transgene insertion in CVH loci and the DNA sequences of amplicons. (E) Immunostaining of PGC-specific markers and mCherry in gSAMURAI PGCs (scale bar: 50 μm).

We next verified the cellular characteristics and potency of gSAMURAI PGCs. Immunofluorescence revealed that PGC-specific markers such as DAZL, DDX4, and NANOG were positively stained in gSAMURAI PGCs (Fig. 3E). These results suggested that the characteristics of PGCs were maintained after genetic manipulation and propagation.

### Generation of gSAMURAI chimeric chickens

The gSAMURAI germline chimeric chicken was generated by transferring PGCs carrying the gSAMURAI transgene into the blood vessel of the embryonic Day 3 (E3) embryo (Fig. 4A). Parts of treated embryos were incubated until E16 and sacrificed to confirm the gonadal colonization of gSAMURAI PGCs. In the gonads from both male and female recipients, the mCherry-expressing cell population could be obviously found (Fig. 4B), suggesting that gSAMURAI PGCs retained the ability to migrate via circulation and colonize the gonads. In the ovary of a 150-day-old hen that was transplanted with gSAMURAI PGCs, several follicles were found to present mCherry fluorescence (Fig. 4C). Semen samples collected from the PGC-transplanted chimeric (F0) roosters were examined for the PCR amplification of transgene fragments. In Fig. 4D, the PCR fragment specific for the CVH KI allele was generated in the DNA samples from gSAMURAI PGCs and F0 rooster semen but not in those from WT rooster semen, confirming that the KI transgene cassette was placed in the CVH locus as designed in the population of gSAMURAI PGCs and F0 rooster sperm.

**Fig. 4.**
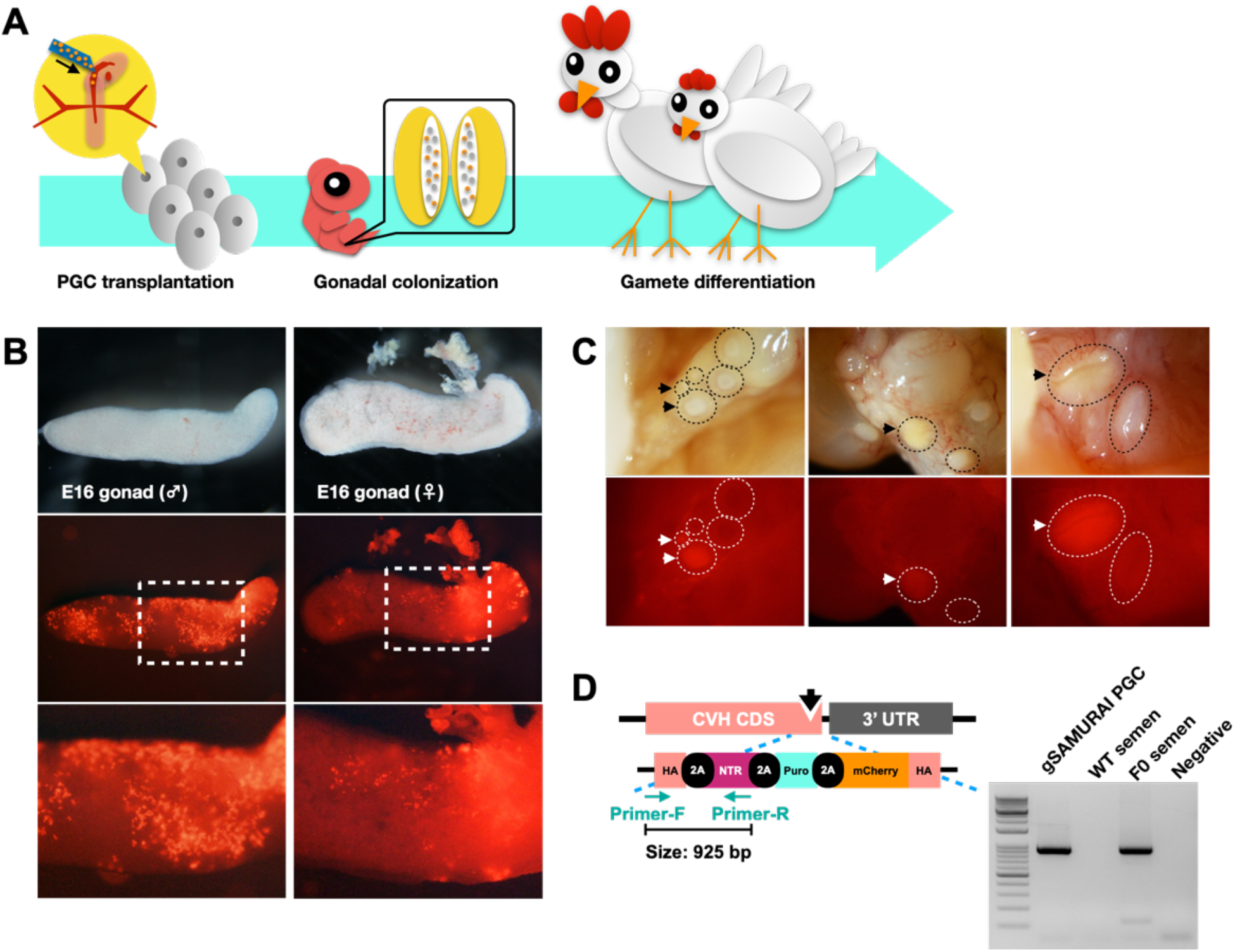
Characterization and function assay in CVH homologous KI - gSAMURAI PGCs. (A) Flowchart of transplantation as the functional assay for gSAMURAI PGCs. (B) The chick gonads at E16 of incubation were isolated from the PGC-transplanted embryos to visualize mCherry fluorescence on transplanted cells. (C) Follicles of the ovary dissected from PGC-transplanted F0 hens showed mCherry fluorescence (arrows). (D) PCR amplification of the DNA fragment (925 bp) crossing the endogenous CVH CDS to the transgene region in the DNA samples from gSAMURAI PGCs, WT rooster semen, and PGC-transplanted F0 rooster semen. DNA-free reaction is shown as a negative control.

### gSAMURAI offspring presented germline-specific mCherry expression visualizing germ cells

We generated gSAMURAI offspring by intercrossing the germline chimeric rooster (F0) to WT hens with artificial insemination. To confirm the inheritance of the gSAMURAI transgene, F1 embryos were dissected at E12 (Fig. 5A). Seven out of 37 (18.9%) embryos carried the gSAMURAI transgene, and these embryos presented mCherry expression in germ cells of both site gonads in both sexes, while comparatively few mCherry-expressing cells were found in the right site gonad compared to the left in the female. This is consistent with the nature of asymmetry between the left-right gonad in gonadogenesis; the left site shows a great number of germ cells, especially in the female gonad (Intarapat and Stern, 2013). We isolated both sites of gonads from WT and gSAMURAI embryos (Fig. 5B). No fluorescent cells were found in the WT gonads regardless of sex. On the other hand, gSAMURAI embryonic gonads showed robust mCherry expression in both site gonads of the male (CVH^KI/Z^) and the dominant site gonad (right) of the female (CVH^KI/W^).

**Fig. 5.**
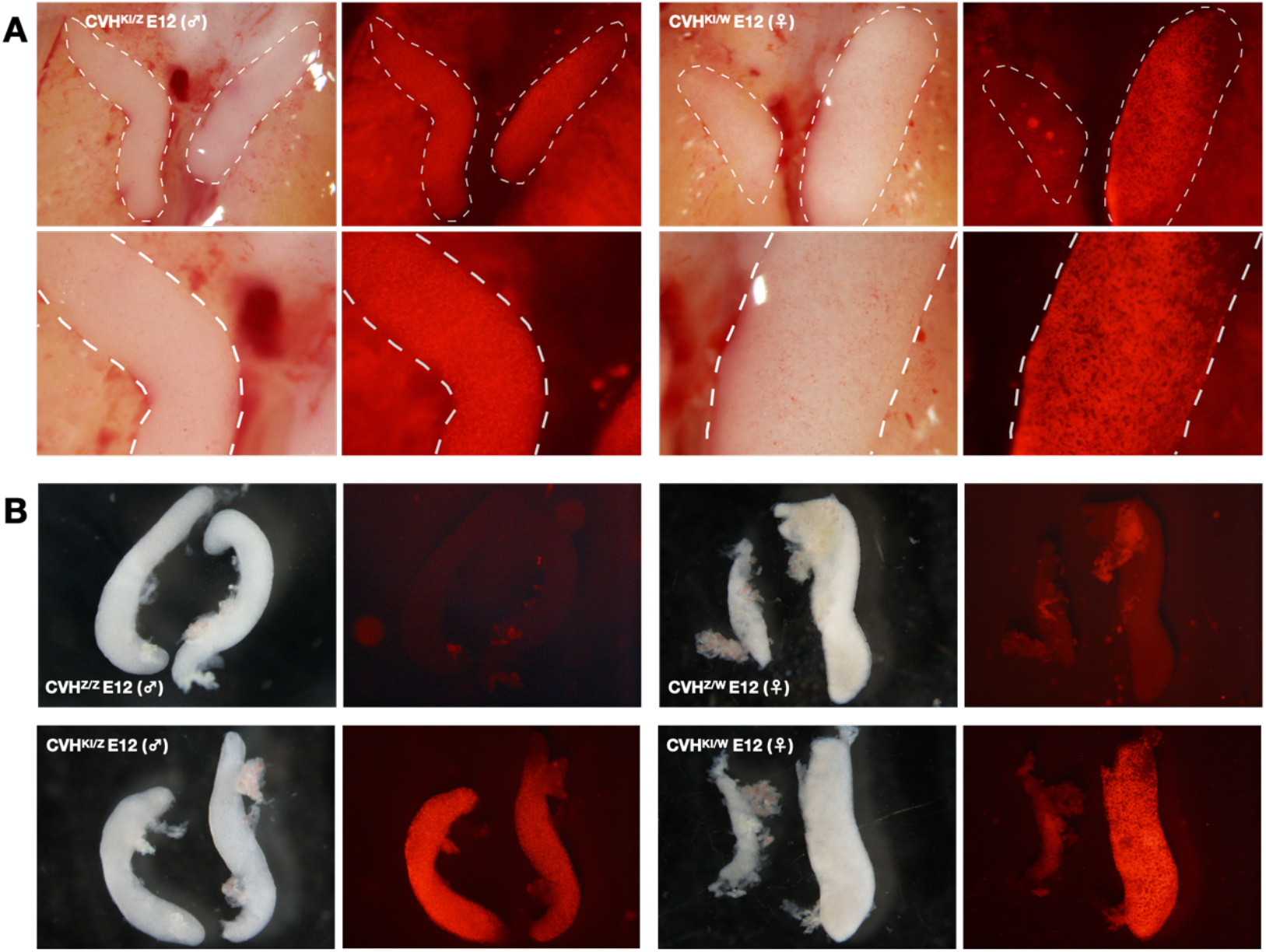
Germline-specific mCherry expression in gSAMURAI chick embryonic gonads. (A) Heterozygous gSAMURAI E12 males (CVH^KI/Z^) and females (CVH^KI/W^) showed mCherry expression in germ cells of gonads (white dotted line). (B) Dissociated gonads from WT chick embryos presented no fluorescence that reflected brilliant mCherry expression in gonads isolated from gSAMURAI chick embryos.

We also observed the fluorescent germ cells of gSAMURAI embryos at E2 with an ex ovo embryo culture. In Movie 1, gSAMURAI germ cells expressing mCherry were circulating in the blood vessels at E2.5 (HH13-14), and a small proportion of them arrested at the area between two bilateral dorsal aortae to micro vessels and near the region for genital ridge. This phenomenon continued at the E3 stage. Movies 2 & 3 showed germ cell efflux from the dorsal aorta and stopped in small capillaries. The area with arrested germ cells was enlarged to cover almost all the outer sides of the bilateral dorsal aortae behind the tail bud and condensed in the region of the genital ridge (Movie 4) (Saito et al., 2022).

These results demonstrated that the genome-edited gSAMURAI PGCs could be inherited by the next generations. Additionally, the resulting gSAMURAI offspring showed germ cell-specific fluorescence, enabling the tracking of germ cells in live embryos.

Movie 1. The germ cell circulation at the region surrounding the genital ridge of an ex-ovo cultured E2.5 gSAMURAI embryo.

Movie 2. The germ cell circulation at the region surrounding the genital ridge of an ex-ovo cultured E3 gSAMURAI embryo.

Movie 3. Migrating germ cells arrested at small capillaries when effluxed from the dorsal aorta to capillaries in E3 gSAMURAI embryos.

Movie 4. An enlarged view and arrow labels migrating germ cells arrested at small capillaries in E3 gSAMURAI embryos.

### Ablation of gSAMURAI PGCs by administering Mtz enriches exogenously transplanted PGCs

We examined whether inducible germline-specific cell ablation works in gSAMURAI chickens. First, we administered a prodrug of the NTR/Mtz system, Mtz, when transplanting exogenous PGCs into gSAMURAI embryos. To distinguish the exogenous PGCs from gSAMURAI PGCs, PGCs expressing mClover under the control of chicken NANOG, a PGC marker gene, were utilized as donor PGCs (Fig. S1). As shown in Fig. 6A, we transplanted NANOG-mClover PGCs into gSAMURAI E3 embryos together with different dosages of Mtz and incubated these embryos until E10. Fig. 6B shows the colonization of gSAMURAI and NANOG-mClover PGCs in gonads. In the non-prodrug control group, both transplanted NANOG-mClover PGCs and host gSAMURAI PGCs shown by mCherry expression were found in the gonads in a well-distributed manner. On the other hand, in the Mtz administration groups, the occupancy of gSAMURAI PGCs in gonads was obviously reduced, whereas transplanted NANOG-mClover PGCs were well distributed. In the quantification results (Fig. 6C), the transplanted PGC rate in the total recipients of the Mtz administration groups was significantly higher than that in the control group (*P*<0.05). Furthermore, the quantified data were scattered by the sex of recipients. Female recipients in the Mtz administration groups showed higher average exogenous PGC rates than those in the control group, but no significant difference was found. In contrast, among male recipients, the exogenous PGC rate in both doses of Mtz administration groups was significantly higher than that in the control group (*P*<0.05). Moreover, all higher exogenous PGC rates (>70%) in the Mtz administration groups were found in males. The highest exogenous PGC rate was 28.77% in the control group, and the highest rates were 79.67% and 90.15% in the Mtz-1 mM and Mtz-5 mM groups, respectively.

**Fig. 6.**
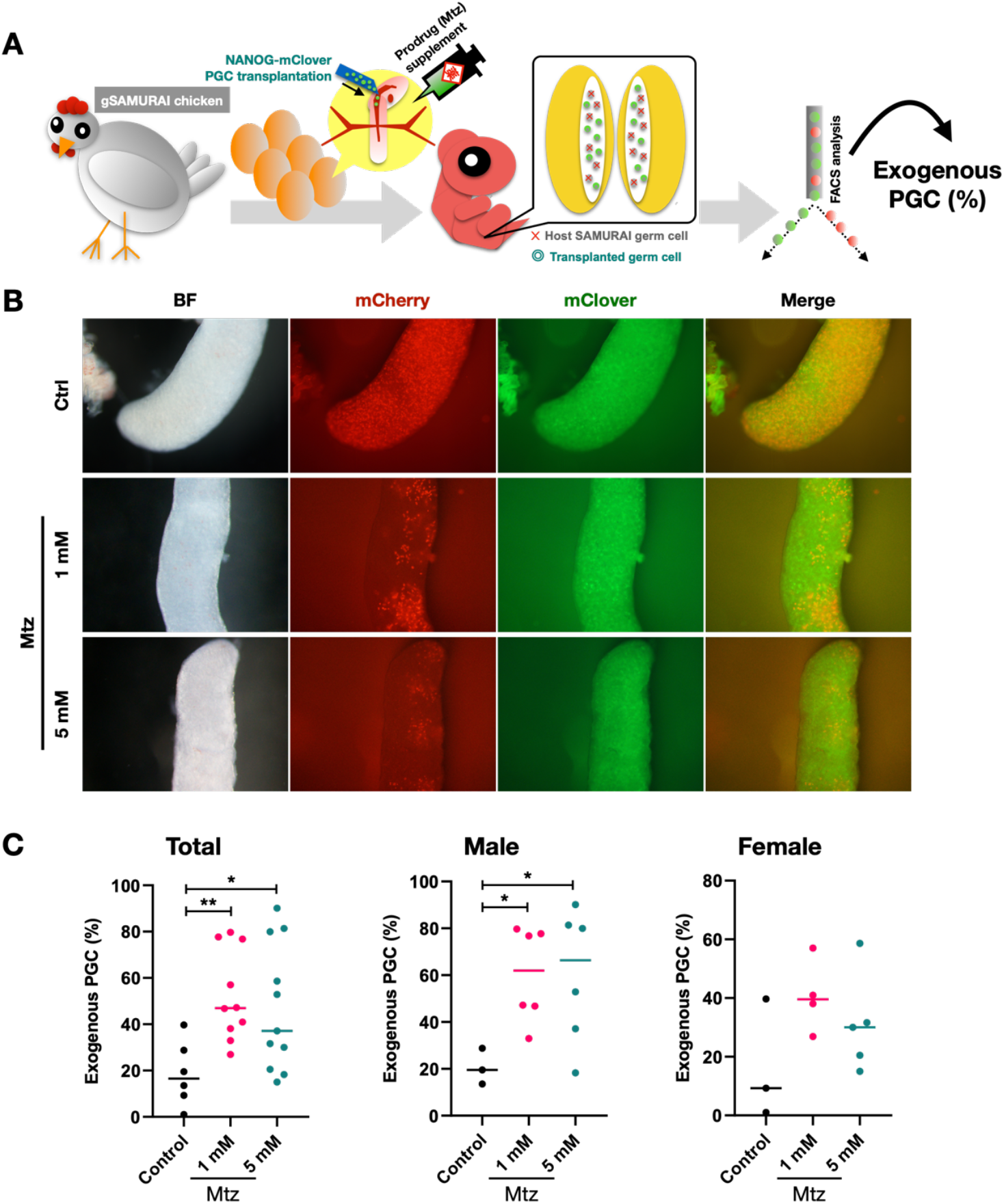
Prodrug-Mtz supplementation with PGC transplantation in gSAMURAI recipients. (A) Schematic of the Mtz administration test using gSAMURAI chick embryos as a recipient to compare the rate of exogenous PGC after PGC transplantation. (B) E10 gonads dissociated from NANOG-mClover PGC-transplanted gSAMURAI embryos with different treatments. (C) Quantification of the exogenous PGC rate derived by FACS analysis in different treatment groups (^*^*P* value of <0.05; ^**^*P* value of <0.01).

These results indicated that the NTR/Mtz system functions to ablate the host germ cells in gSAMURAI embryos without affecting exogenous PGC migration and colonization. An extreme predominance in the donor germ cell population was generated by using the gSAMURAI chicken model as the recipient, indicating a largely increasing potential to obtain the target offspring after sex maturity.

## Discussion

Recently, several studies reported the production of gene-edited chickens using transcription activator-like effector nucleases (TALENs) or the CRISPR-Cas9-mediated method (Oishi *et al*., 2016; Park et al., 2014). In this study, we successfully generated a genetically modified chicken with the CRISPR-Cas12a (Cpf1) family nuclease MAD7. MAD7 differs from Cas9 in its nuclease recognition sequence and the DNA end sequence after cleavage. MAD7 recognizes a thymidine-rich sequence in the PAM site upstream of the crRNA, whereas Cas9 uses a guanine-rich PAM sequence downstream of the crRNA. crRNAs of Cpf1 family nucleases, including MAD7, are shorter than Cas9 single guide RNAs (sgRNAs) due to the differences in repeat- and tracrRNA-derived segments. The Cpf1 family crRNA is only ∼43 nt in length, while the most commonly used sgRNA scaffold for Cas9 is ∼101 nt in length (Swarts and Jinek, 2018). Concerning the DNA end sequence after cleavage, MAD7 uses a single RuvC-like endonuclease domain to cut each DNA strand to form a sticky end, whereas Cas9 uses HNH and RuvC nuclease domains to form a blunt-ended DSB, and (Swarts and Jinek, 2018). The two different types of gene editing nucleases allow for a wider range of target designs.

In the present study, we demonstrated that MAD7 introduced a mutation into the CVH loci of chicken PGCs at ∼10% indel frequency. Compared to the indel formation from 4% to 23% found in a human cell line when MAD7 was used, it was in the range of prediction (Liu et al., 2020). By using this nuclease for DNA cleavage-mediated homologous recombination, gene KI in the CVH loci of chicken PGCs was obtained. Following the selection of an antibiotic for 2 weeks, almost a hundred percent of KI PGCs could be achieved without affecting the PGC characteristics. MAD7 shows sufficient availability in the genetic modification among chicken PGCs as other nucleases.

We generated gSAMURAI chickens expressing a germline-specific mCherry reporter, which will be a powerful tool in the study of germ cell development. CVH, also known as DEAD-Box Helicase 4 (DDX4), was used as the leading gene for the reporter. CVH/DDX4 is a metazoan-conserved protein that encodes an RNA binding protein and plays an important role in germline formation (Mochizuki et al., 2001). Its protein expression was detected in the embryo since the first cleavage stage, which localized in the cytoplasm of germ cells covering PGCs in the embryo to gametes in adult gonads (Tsunekawa et al., 2000). Avian species are known to translocate PGCs from the extraembryonic region to gonads through circulation after being enveloped in vascular tissue, which is different from the mammalian case (Murai et al., 2021; Nakamura et al., 2013). In the present study, using an ex ovo culture of E2.5-E3 gSAMURAI embryos, we detected mCherry-expressing PGCs circulating in and exiting from blood vessels. PGCs were found to migrate within a regular pattern such that a pair of the mainstream dorsal aorta accelerated by a heart pump carried PGCs to enter the tributary stream – the capillary plexus around the genital ridge (Movie 1-4). In this period, PGCs seemed to slow down the velocity even though some of them became stuck in the capillary and arrested the flow. PGC occlusion in Ex-VaP is the critical step for further transmigration into the gonad that was recently determined by our research partners. Saito *et al*. (2022) suggested that actin-mediated regulation of cellular stiffness is highly dynamic in avian PGCs during migration. This extravasation event in avian PGCs could be linked to cancer metastasis since they shared a similar cellular migration manner (Reymond et al., 2013). All of these results indicate a high availability of our gSAMURAI chicken embryo for these investigations since PGC becomes easy to track in this model.

In this study, we demonstrated the efficient ablation of host germ cells using the NTR/Mtz system in a gSAMURAI chicken model. NTR is able to undergo electron transfer from NADH or NADPH to induce a nitro reduction reaction, and cytotoxic alkylating agents are generated when reacting with Mtz, which can cause cell death via the caspase-3-mediated apoptosis pathway (Chen *et al*., 2011). The composition of gonadal occupancy between the endogenous and exogenous germ cells had a dramatic change with a treatment of prodrug, endogenous ones came to the minority population, exogenous ones displayed the dominant roles and obtained approximately 90% as the highest occupancy among samples. This result indicated that the gSAMURAI model can be an ideal recipient of germ cell transplantation for germplasm management and/or new strain development in terms of saving time and cost. According to the sensitivity to the NTR/Mtz system, prodrug administration at a higher concentration (∼10 mM) and enough exposure time (24 hours) were required to deplete NTR-expressing specific cell types in zebrafish (Curado et al., 2007; Hu et al., 2010). However, the administration method was completely different between the fish and chicken models. In our study, the prodrug was directly injected into embryonic circulation instead of being exposed to the living environment, as in fish models. Recently, the Mtz chemical analogs furazolidone and ronidazole were found to have lower working concentrations with NTR, especially in ronidazole, showing better potential as a replacement for Mtz due to its lower nonspecific toxicity (Lai et al., 2021). Therefore, for a more consistent result with high efficiency to deplete targeting cells, the manipulation of prodrug administration in type, method, and exposure period still needs further investigation. The gSAMURAI chicken model has advantages compared to classical methods. In classical strategies, host-derived germ cells have been partially removed by chemical or irradiation treatment, but these strategies showed high embryonic mortality and/or abnormality (Nakamura *et al*., 2012; Nakamura *et al*., 2010). Although our gSAMURAI chicken is required to administer the pro-drug Mtz to remove germ cells, based on our observations, it does not affect the survival rate and embryo development. Recently, genetic methods for this purpose were developed to overcome this problem. CVH null chickens were generated to show germ cell ablation and thus provided an ideal surrogate host for the generation of avian species with PGC transfer (Taylor et al., 2017). Within this genetically sterile model, an almost perfect transmission rate was obtained in a female recipient with the same sexual PGC lines as donors, while the WT recipient female with the identical set of PGC transfers showed no heritage from donor PGCs in the study (Woodcock *et al*., 2019). Nevertheless, it is supposed that the maintenance and production of CVH null chickens need to expend much effort since sterility is a double-edged sword. Therefore, an inducible sterile model is likely more suitable since the side-effect problem in the previous method could be overcome. Ballantyne et al. (2021) described a genetically modified chicken model produced with a gene insertion to express an inducible caspase-9 (iCaspase9) protein led by the endogenous DAZL gene; thus, a switchable germ cell-deficient model was formed. Using this system as a surrogate, they produced chicken offspring carrying edited alleles inherited from donor PGCs with a high germline transmission rate since almost all host germ cells were ablated. However, the iCaspase9 chicken model presents a comparatively low hatchability (∼60%), indicating an influence of the iCaspase9 transgene on chicken embryonic development (Ballantyne *et al*., 2021).

In conclusion, we generated a genetically modified chicken model, namely gSAMURAI, providing not only convenience in germ cell tracking as a tool for research on germ cell development but also as an ideal recipient for every production purpose through germ cell transfer because endogenous germ cells could be depleted with a simple induction process. This function will facilitate production efficiency and thus will be expected to contribute to avian germplasm management for the regeneration of endangered species and the production of novel poultry strains with gene editing techniques.

## Acknowledgments

The authors thank all the members of Setsuro Tech Inc. and of the Laboratory of Embryology, Tokushima University, for their help and kind discussion. We also appreciate all the assistance for animal production and maintenance by Kyodoken Institute.

